# A self-adaptive and versatile tool for eliminating multiple undesirable variations from transcriptome

**DOI:** 10.1101/2024.02.04.578839

**Authors:** Mengji Zhang, Lei Yan, Xinbo Wang, Yi Yuan, Shimin Zou, Sichao Yao, Xinyu Wang, Tian Xu, Bin Chen, Dong Yang

## Abstract

Accurate identification of true biological signals from diverse undesirable variations in large-scale transcriptomes is essential for downstream discoveries. Herein, we develop a universal deep neural network, called DeepAdapter, to eliminate various undesirable variations from transcriptomic data. The innovation of our approach lies in automatic learning of the corresponding denoising strategies to adapt to different situations. The data-driven strategies are flexible and highly attuned to the transcriptomic data that requires denoising, yielding significant improvement in reducing undesirable variation originating from batches, sequencing platforms, and bio-samples with varied purity beyond manually designed schemes. Comprehensive evaluations across multiple batches, different RNA measurement technologies and heterogeneous bio-samples demonstrate that DeepAdapter can robustly correct diverse undesirable variations and accurately preserve biological signals. Our findings indicate that DeepAdapter can act as a versatile tool for the comprehensive denoising of the large and heterogeneous transcriptome across a wide variety of application scenarios.

## Introduction

Transcriptome reveals the diversified cell types, cellular states, and regulatory mechanisms of molecular underpinnings by quantifying the dynamically transcribed RNA^1^, serving as a crucial tool to decipher the biological mechanisms. With rapid development^1-3^ and widespread applications of transcriptome^4-6^, enormous transcriptomic data has been accumulated over the past two decades^7-9^. These heterogeneous datasets not only provide invaluable resources but also present new computational challenges in delivering comprehensive biological insights from integrated analyses due to the presence of diversified unwanted variations.

These undesirable variations fall into four categories: variabilities from notorious batch effects (batch variation)^10^, sequencing platforms (platform variation)^11, 12^, heterogeneous bio-samples (purity variation)^13, 14^ and other unknown technical differences (unknown variation), substantially compromising downstream discoveries. Batch variations arising in different runs at different time points represent the prevailing technical factors^15^. Platform variations include the systematical experimental bias intra- and inter-platforms. Intra-platform variations are brought by varied sequencing depth over orders of magnitude^11, 16^, while inter-platform variations originate from different sequencing technologies, e.g., RNA-seq and microarray^17^. Purity variations refer to the bias among profiles sequenced from heterogeneous bio-samples with diversified cellular composition, e.g., cancer cell lines and tumor tissues^13, 14^. In addition to these three well-defined types of variations, there exist a large number of unknown variations that are obscured and difficult to be manually corrected, confounding the true biological signals. More challenging is that these undesirable variations often coexist, making the quest for true biological discoveries more complicated.

Existing methods aim to remove one of these variations with nearly manually designed strategies to constrained situations for integrated transcriptomic analysis. Batch removal methods hypothesize that the true biological signals and non-biological noises exhibit either a linear^18, 19^ or orthogonal^10, 20^ relationship and significantly mitigate the batch variations by subtracting the estimated batch noises. Still, the rigid assumption falls short in correcting the large and complex transcriptomic datasets featuring co-existing non-orthogonal variations, particularly in the era of big data. Normalization methods are designed to eliminate the intra-platform variations from profiles sequenced by the same platform with the implicit assumption that transcriptomic readouts are proportional to some scaling factors^21, 22^. Correspondingly, variation removal is implemented by simply dividing all gene readouts in one sample using a defined scale factor^23, 24^. The challenge emerges when datasets come from different platforms, namely the inter-systematical variations, making the inter-platform gene readouts cannot be corrected with one simple scale factor. For purity variations, bulk deconvolution methods successfully estimate the relative cellular abundances from heterogeneous bio-samples^25^. Yet, deconvolution methods fail to reconstruct the denoised transcriptomic spectra, showing limitations in correcting composition differences across bio-samples. As for unknown variations, manual schemes fail to tackle this kind of variation due to the high dependence on pre-defined assumptions.

Herein, we develop a deep neural network, called DeepAdapter (**Deep A**lign large and **d**iversified transcriptome with **a**dversarial network and **d**eep me**t**ric l**e**a**r**ning), to robustly remove undesirable variations from large and heterogeneous transcriptomes. Tailored to adapt to various application scenarios automatically, DeepAdapter learns the latent space where distances of bio-samples that are collected from diverse data sources yet should carry similar biological signals are minimized. Especially, the fact that DeepAdapter requires no prior information endows it a great advantage in eliminating unknown variations. These flexibilities facilitate the automatic learning of diverse denoising strategies, making it suitable for various application scenarios.

We evaluated DeepAdapter thoroughly on multiple tasks, and it considerably outperforms state-of-the-art methods in variations correction and biological signal conservation. For batch variations, by exploiting transcriptomic datasets derived from multiple batches, we reveal that DeepAdapter can eliminate the conventional batch variations and conserve the distinct donor-wise signals. For platform variations, we utilize large RNA-seq and microarray profiles of the same cancer cells and demonstrate that DeepAdapter can remove these variations and advance cancer subtype identification across different platforms. For purity variations, by employing RNA-seq profiles of cancer cell lines and tumor tissues, we illustrate that DeepAdapter can correct variations, including but not limited to those caused by varied tumor purity and immune infiltration, thereby enhancing lineage identification and reproducing the associations between prognostic marker gene expression and clinical survival outcomes across heterogeneous bio-samples. For unknown/mixed variations, we conceptualize the blind information of other technical factors, such as laboratory sites and library preparation protocols, as the unknown variations and demonstrate that DeepAdapter can eliminate these undesirable variations and recover true gene co-expression signals.

## Results

### Datasets with undesirable variations

#### Transcriptomic data with batch variations

We include 2 set of transcriptomic datasets to study the batch variations: a) Batch-LINCS: RNA-seq profiles of cardiomyocyte-like cell lines across 4 batches are sourced from the LINCS DToxS project^26^. The transcriptomic profiles are seque nced from 706 samples within 4 donors. The gene set is the intersection of 10112 transcripts across 4 batches. b) Batch-Quartet: RNA-seq profiles of B-lymphoblastoid cell lines spanning 21 batches are sourced from Quartet project^27^. The profiles are collected from 252 samples within 4 donors. The gene set is the intersection of 58395 transcripts across 21 batches.

#### Transcriptomic data with platform variations

We include cell lines sequenced by RNA-seq and microarray technologies to study the variations brought by technology revolutions. Microarray of cell lines is sourced from The Genomics of Drug Sensitivity in Cancer (GDSC) project^8^ that covers 948 samples spanning 30 cancer types. RNA-seq of cell lines is sourced from CCLE (22Q2 project)^9^ that covers 1406 samples within 32 cancer types. The gene set is the intersection of 16308 transcripts between microarray and RNA-seq. The curated dataset is referred to Platform-GDSC-CCLE.

#### Transcriptomic data with purity variations

We include RNA-seq data of cell lines and tumor tissues to study the variations caused by heterogeneity in different bio-samples. RNA-seq of cell lines comes from CCLE (19Q4 project)^9^, which covers 1249 samples within 32 cancer types, and RNA-seq of tumor tissues comes from The Cancer Genome Atlas (TCGA) project^7^, which covers 9806 samples spanning 26 cancer types. The transcriptomic data covers 36631 protein-coding genes derived from the intersection between cell lines and tumor tissues. The curated dataset is referred to Purity-TCGA-CCLE.

#### Transcriptomic data with unknown/mixed variations

In this work, we utilize Batch-Quartet and TCGA datasets to illustrate the unknown/mixed variations. For Batch-Quartet, DeepAdapter is blinded to the information of library preparation protocols, RNA-seq devices, and laboratory sites, simulating these variations as unknowns. For TCGA, we refer to the information of different processing dates and batches as the unknown/mixed variations in the same manner.

### Overview of DeepAdapter

DeepAdapter is designed to remove a wide range of different variations in transcriptomics data, thereby enabling meaningful biological discoveries with aligned large-scale datasets. These variations can be brought by batch effects^15^, sequencing technology revolutions (e.g., from microarray to RNA-seq)^28^, inherent heterogeneity of bio-samples (e.g., cell lines and tumor tissues)^14^, etc. To achieve this goal, we design DeepAdapter, independent of any prior distribution assumptions, serving as a universal adapter for integrating large and heterogeneous transcriptomes (Figure 1a).

**Figure 1.**
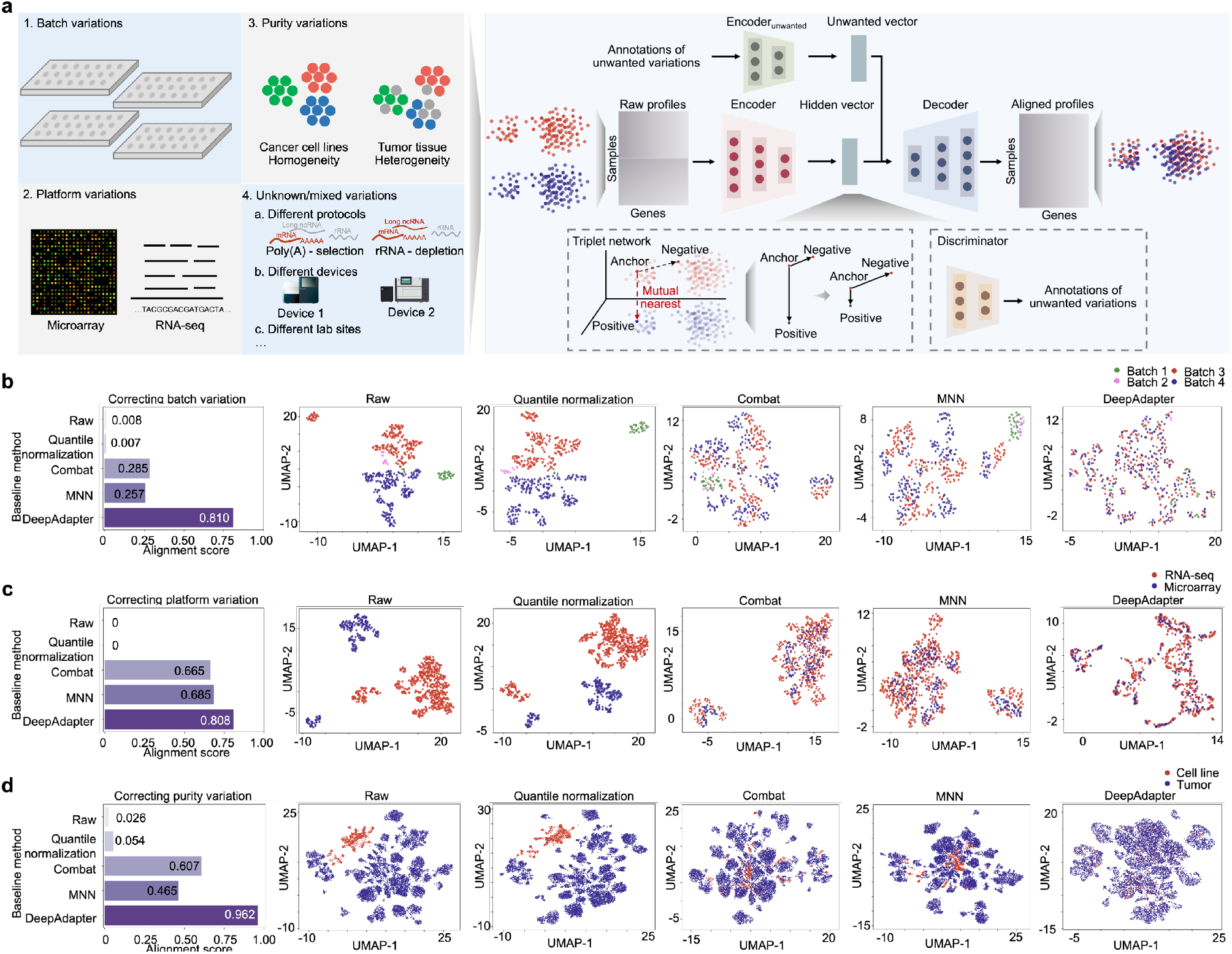
Overview and performance assessment of DeepAdapter. a) Diverse sources of unwanted variations in transcriptome and the architecture of DeepAdapter. DeepAdapter is developed in an autoencoder, where the variations removal is performed in the latent space between the encoder and the decoder. Specifically, a discriminator and a triplet network are trained to remove the unwanted variations. b) The alignment performances of different methods across batches (batch variations). c) The alignment performances of different methods across sequencing microarray and RNA-seq (platform variations). d) The alignment performances of different methods across tumor cells and tumor tissues (purity variations).

#### Structure of DeepAdapter

We construct a nonlinear autoencoder to extract profiles of true signals and remove the unwanted variations in the latent space. Specifically, we design a discriminator and a triplet neural network to remove these variations. Inspired by adversarial learning^29^, the discriminator is trained to classify the annotations of undesirable variations using the learned latent vectors. Additionally, distances of bio-samples that carry similar biological information but are collected from different data sources should be minimized. Therefore, we firstly define anchor, positive, and negative samples: anchor samples are enumerated from the training set, positive samples are defined as the mutual nearest neighbors (MNN)^10^ between data sources and negative samples are randomly sampled from the anchor dataset. Then, we utilize triplet network to minimize the distance between anchor and positive samples while maximizing that between anchor and negative samples (Methods).

#### Baseline comparison of alignment performance

We compare the performances on variation removal with quantile normalization (QuanNorm)^30^, Combat^18^, and MNN^10^. We utilize the UMAP^31^ to visualize the results and alignment score^32^ to assess the quantitative performances (Figure 1b-d). The raw data is greatly affected by the undesirable variations described above, with an alignment score of 0∼0.026 (Figure 1b-d). Strikingly, DeepAdapter dramatically eliminates these variations and exhibits a significant improvement in alignment performance compared to baseline methods: 1) alignment score of 0.810/0.285/0.257/0.007 for DeepAdapter/Combat/MNN/QuanNorm across batches, 2) alignment score of 0.808/0.685/0.665/0 for DeepAdapter/MNN/Combat/QuanNorm across sequencing platforms, and 3) alignment score of 0.923/0.607/0.465/0.054 for DeepAdapter/Combat/MNN/QuanNorm across bio-samples with different purities.

Next, we dive deeper into the correction results from our model in three different scenarios described above. With these examples, we demonstrate the effectiveness of DeepAdapter in removing diverse variations and preserving biological signals for large and heterogeneous transcriptomic datasets.

### Correction the batch variations

Batch effects are the common challenge in transcriptomic investigations, stemming from technical factors like instrumental calibration and reagent variability. Remarkably, these variations often obscure true biological signals, introducing complexity and confounding subsequent analyses.

#### DeepAdapter eliminates the batch variations

To assess the performance in correcting batch variations, we employ DeepAdapter to correct the batch effects in Batch-LINCS and Batch-Quartet datasets. Obviously, raw transcriptomic data shows clear separations among batches (Figure 2b&e) with poor alignment score (0.307/0.011) and average silhouette width (ASW)^20^ score (0.462/0.326) for Batch-Quartet and Batch-LINCS (Figure 2a&d). With DeepAdapter, the evaluation metrics have been largely improved with alignment score of 0.731/0.856 and ASW score of 0.572/0.543 across Batch-Quartet (4 batches, Figure 2a-b) and Batch-LINCS (21 batches, Figure 2d-e), respectively.

**Figure 2.**
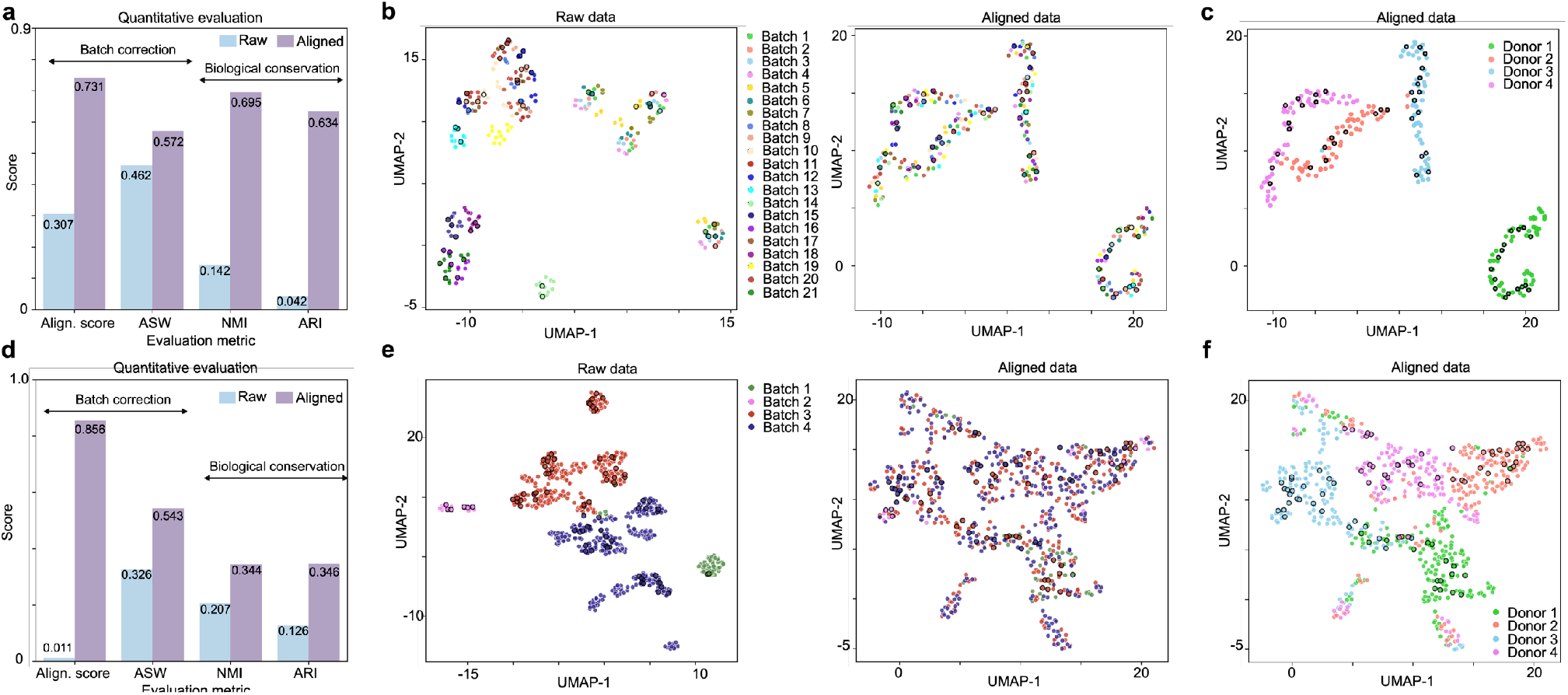
Alignment of datasets profiled across batches. a) The quantitative metrics of batch correction and biological signals conservation for Batch-Quartet data. b) The UMAP analysis of raw and aligned data across 21 batches for Batch-Quartet data. c) The UMAP analysis of aligned data across 4 donors for Batch-Quartet data. d) The quantitative metrics of batch correction and biological signals conservation for Batch-LINCS data. e) The UMAP analysis of raw and aligned data across 4 batches for Batch-LINCS data. f) The UMAP analysis of aligned data across 4 donors for Batch-LINCS data. White-edged and black-edged circles refer to the training and testing set, respectively.

#### DeepAdapter conserves the distinct donor-wise signals

Donor sources are representative biological signals. We observed DeepAdapter clearly separate samples from distinct donors into clusters, showing a satisfactory performance of biological signals conservation. The results were further supported by the normalized mutual information (NMI)^33^ and adjusted rand index (ARI)^34^ analyses. Specifically, DeepAdapter makes a noteworthy improvement in NMI score, progressing from 0.142/0.207 to 0.695/0.344, and ARI score, advancing from 0.042/0.126 to 0.634/0.346, for Batch-Quartet (Figure 2c & Figure S2) and Batch-LINCS (Figure 2f & Figure S2). These findings illustrate the robust efficacy of DeepAdapter in accurately preserving biological signals across batches.

### Correcting the platform variations

Sequencing technologies for transcriptomics have evolved rapidly from microarray to RNA-seq. Integrated analyses of datasets from these two technologies remain challenging due to their inherent variability difference. Microarray platforms detect the signals with sophisticated probe sets, while RNA-seq technologies record the counts of copy numbers for transcripts. Consequently, microarray measures the continuous values of transcripts following the normal Gaussian distribution, and RNA-seq records the integer counts, thereby making the expression profiling incomparable and hampering meaningful biological discoveries.

#### DeepAdapter eliminates the platform variations

To evaluate the performance of DeepAdapter in eliminating platform variations, we use DeepAdapter to remove the nonbiological effects between microarray and RNA-seq datasets and generate an integrated gene expression dataset across sequencing platforms. As expected, raw data shows an obvious separation between microarray and RNA-seq technologies with a poor alignment score of 0 and ASW score of 0.265 (Figure 3a-b), consistent with the differential distributions in gene expression (Figure 3d). Correspondingly, DeepAdapter substantially improves the alignment/ASW score to 0.833/0.496 (Figure 3a&c) and reduces the divergence score from 226.14 to 2.41 across microarray and RAN-seq technologies (Figure 3d).

**Figure 3.**
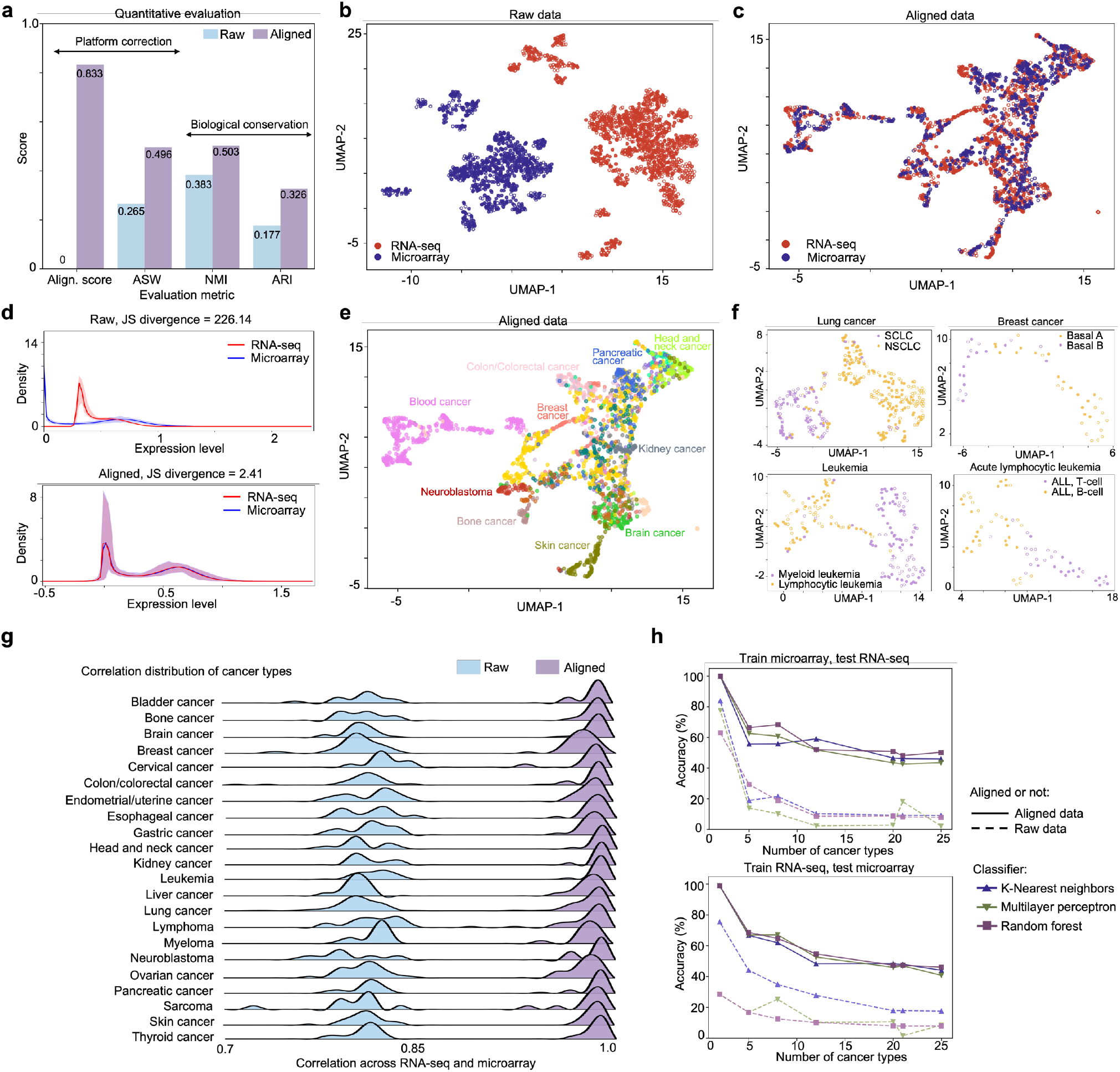
DeepAdapter enables the correction of the platform variations. a) The quantitative metrics of platform correction and biological signals conservation. The UMAP analysis of b) raw and c) aligned data across microarray and RNA-seq platforms. d) Divergence score of raw and aligned data. e) The UMAP analysis of aligned data across cancer types. f) The UMAP analysis of aligned data for molecular subtypes. SCLC refers to small cell lung cancer. g) Correlation distribution of the identical tumor cells profiled by microarray and RNA-seq across cancer types. h) Cross-sequencing-platform classification performances of raw and aligned data using DeepAdapter. Solid and dashed lines refer to the performances of aligned and raw data, respectively.

#### DeepAdapter preserves meaningful biological relationships across sequencing platforms

To assess the effectiveness of DeepAdapter in preserving biological variations, we first evaluate whether DeepAdapter erased information about known cancer types and subtypes across microarray and RNA-seq datasets during the correction process. As demonstrated in Figure 3e, DeepAdapter corrects much of the systematic variations while reproducing clear cancer-type clusters across different platforms, including lymphoma, myeloma, leukemia, skin cancer, brain cancer, bone cancer, neuroblastoma, colon/colorectal cancer, breast cancer, head and neck cancer, and kidney cancer (Figure S4). Quantitative analyses confirm that our method improves the clustering of tumor types (Figure 3a & Figure S3, NMI/ARI score from 0.383/0.177 to 0.503/0.326). To extend this analysis, we annotate the cancer subtype labels of aligned transcriptomic data for microarray and RNA-seq. The results reveal that molecular subtype information is aligned with DeepAdapter. For example, lung cancer and breast cancer subtypes are aligned in the same tumorous space (Figure 3f). Correspondingly, leukemia cell lines formed two distinct clusters, a myeloid cluster and a lymphocytic cluster (Figure 3f) with B and T cells in distinguishable clusters (Figure 3f).

In addition, we explored the similarity between the corresponding transcriptomes that were measured by both microarray and RNA-seq techniques (n=684). Interestingly, compared to the correlation of 0.798-0.831 in the original data, the correlation has been significantly improved to 0.978-0.993 after removing the platform variation by DeepAdapter (Figure 3g).

A huge number of transcriptomic data are available today through various platforms. However, they are difficult to integrate and, therefore, not fully utilized. For example, modern artificial intelligence and machine learning capabilities rely heavily on the availability of large amounts of high-quality data. We thus assessed the potential to integrate datasets from different sources. To simulate this situation, we design the cross-platform classification experiment, where machine learning models for cancer-type classification are trained on only microarray datasets, and then tested with RNA-seq data strictly. Strikingly, after the correction of undesirable variation by DeepAdapter, machine learning models extensively improve the classification performances. In the experiment of training on microarray datasets and testing using RNA-seq datasets (Figure 3h & Figure S5), the accuracies (0.464/0.590/0.686) for classifying 25/12/5 cancer types significantly increase, compared to limited accuracies (0.106/0.103/0.252) in the unaligned data. The improved accuracies of multi-classification are also observed in the counterpart setting of training on RNA-seq datasets and testing on microarray datasets (Figure 3g).

Taken together, our results indicate that DeepAdapter has successfully preserved transcriptomic features representative of these biological variations.

### Correcting the purity variations

The tumor masses are highly heterogeneous structures composed of both malignant and nonmalignant cells^35^. Due to the heterogenicity of the tumor samples, numerous bulk transcriptomic data accumulated over the past two decades are compromised by the presence of normal cells to varying degrees. Deconvolution methods have successfully demonstrated the feasibility of inferring cell type compositions from bulk RNA-seq integrated with scRNA-seq as prior knowledge^36^. However, extracting the tumor-specific profiles by eliminating the purity variations beyond estimating the cellular composition stands as a significantly challenging but pioneering endeavor.

#### DeepAdapter eliminates purity variations with improved tumor purity

To challenge the performance of DeepAdapter in removing purity variations, we apply DeepAdapter to align the gene expression profiles measured from cancer cell lines and human tumor tissues (Figure 4). The underlying hypothesis is that if transcriptomic differences between tumor cell lines and tumor masses are corrected by DeepAdapter, then the majority of the eliminated signals will be purity variations. To test this hypothesis, we compare the transcriptomic data before and after alignment by DeepAdapter and find that cell lines and tumor tissues are well-aligned with alignment/ASW score (Figure 4a) impressively improved from 0.030/0.285 to 0.961/0.499 (Figure 4b-c). Interestingly, additional analyses validate that this result is largely due to the removal of purity variations. To ensure rigor, we apply two metrics (tumor purity score and immune score)^37^ to assess tumor purity and, more specifically, the proportion of infiltrating stromal and immune cells from transcriptomic data. DeepAdapter remarkably improves the estimated tumor purity (Figure 4g & Figure S6) and decreases the presence of immune cells for 96.9% (31 / 32) of aligned cancer types (Figure 4h-i & Figure S6).

**Figure 4.**
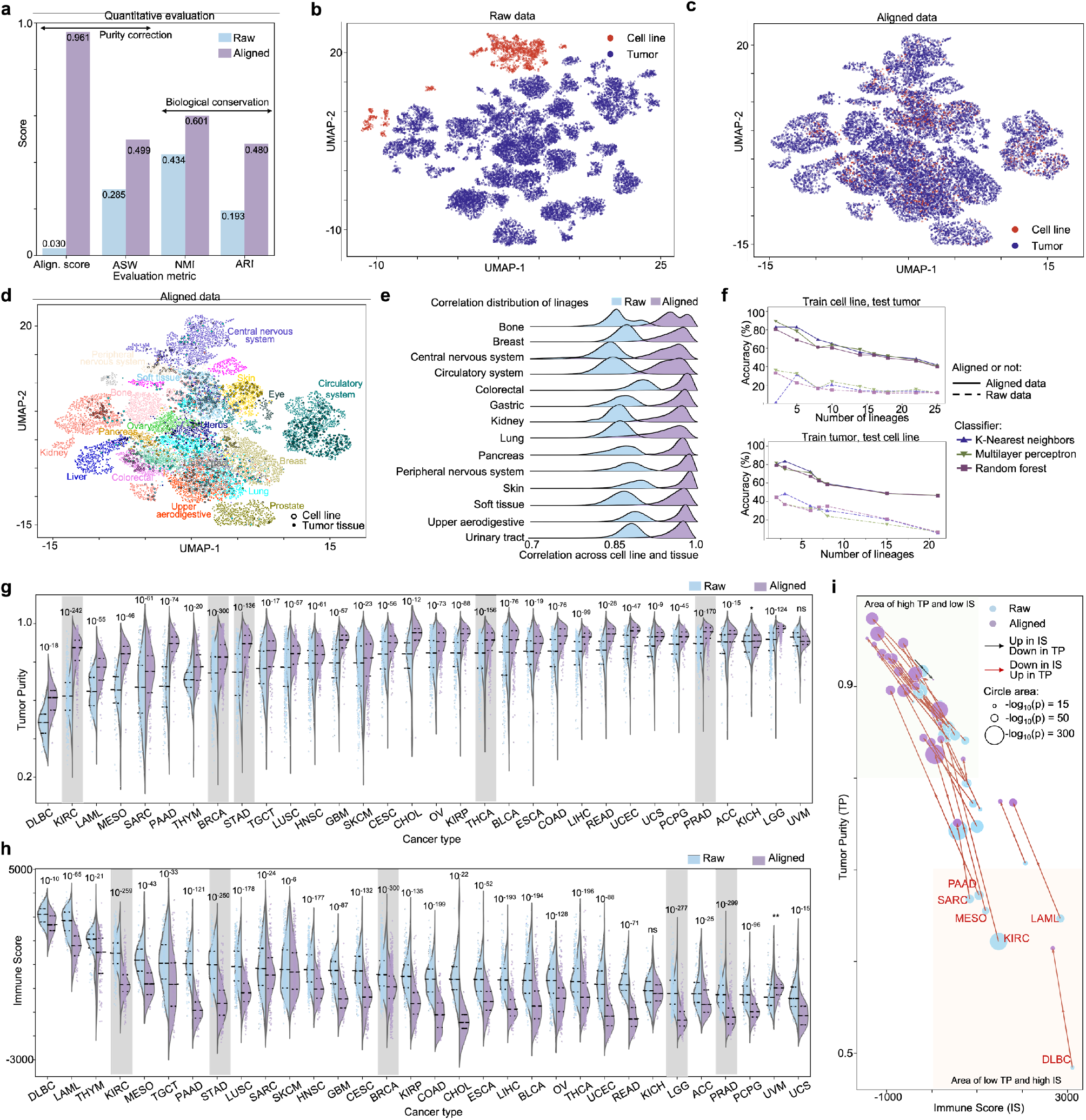
DeepAdapter enables the correction of the purity variations. a) The quantitative metrics of batch correction and biological signals conservation. The UMAP analysis of b) raw and c) aligned data across tumor cells and tumor tissues. d) The UMAP analysis of aligned data across lineages. e) Correlation distribution of profiles sequenced from cancer cell and tumor tissues across cancer types. f) Cross-biosample classification performances of raw and aligned data using DeepAdapter. Solid and dashed lines refer to the performances of aligned and raw data, respectively. g) Analysis of tumor purity for 32 cancer types with aligned data. h) Analysis of immune score for 32 cancer types with aligned data. j) The tumor purity and immune score of individual cancer types. Each dot corresponds to the metrics (immune score (IS), tumor purity (TP)) for each descriptor. The diameter is proportional to the significance level (-log_10_(p)) of the changed TP. The orange region indicates cancers with low TP and high IS. The green region indicates cancers with high TP and low IS.

#### DeepAdapter improves lineage fidelity across bio-samples

To confirm that meaningful biological features are retained, we inspect whether the information about the tumor tissue origin is preserved. As demonstrated in Figure 4d, the corrected gene expression profiling of tumor tissues is clustered with the cell lines of the corresponding tissue origin. Notably, the intra-lineage similarity between cell lines and tissues is improved from 0.872 to 0.964 across lineage types (Figure 4e & Figure S7).

To confirm this observation, we design the cross-biosample classification experiment. we train the cross-biosample classifier with only transcriptomic data of cell lines and test with tumor tissues strictly. DeepAdapter largely improves the accuracy by 37.352% on average. More specifically, DeepAdapter significantly improves accuracies to 0.516/0.696/0.831/0.891 for classifying 18/8/5/2 lineages, compared to 0.158/0.174/0.325/0.360 for the unaligned data (Figure 4f) with a limited number of training samples (169/255/304/274).

#### DeepAdapter reproduces the prognostic marker in survival analysis

Undesirable variations can compromise the downstream identification of prognostic markers from transcriptomic data of patient tumor tissues^11^. For example, SUCLG2P2, GPC1, CISH, and ANKZF1 are identified as survival-associated genes in colon adenocarcinoma (COAD)^11, 38^. However, no such association was detected in unaligned TCGA COAD data (n=262, Figure 5a). Other examples are FBXL14, PKP2 in rectum adenocarcinoma (READ) and STAB1 in breast cancer (BRCA) respectively, whose associations with survival have been previously reported^11, 38^, but these associations were also obscured in uncorrected datasets (Figure 5b-c). Surprisingly, after DeepAdapter correction, we found that these associations were all reproduced (Figure 5).

**Figure 5.**
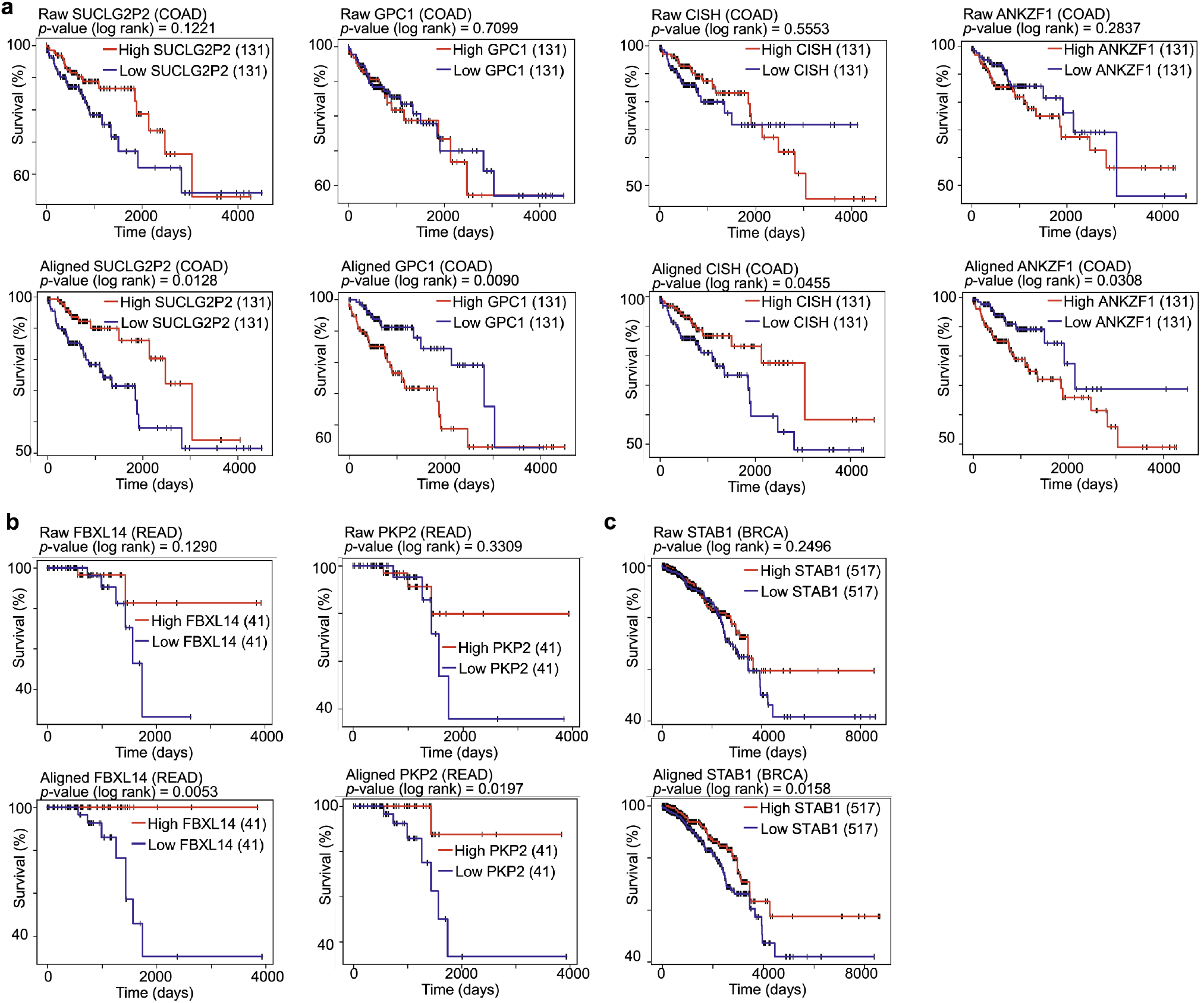
DeepAdapter reproduces the associations of prognostic marker with overall survival. Overall survival analysis of gene expression for a) COAD, b) READ, and c) BRCA. First row: survival analysis of raw transcriptomic data. Second row: survival analysis of aligned transcriptomic data.

### Correcting the unknown/mixed variations

In addition to the three well-known variations mentioned above, independent transcriptome studies inevitably introduce a large number of other unwanted variations from a variety of sources. These variations are difficult to define and eliminate, leading to misleading discoveries.

#### DeepAdapter eliminates unknown variations in laboratories, library protocols, and RNA-seq devices

Batch-Quartet study involves 252 samples within 4 donors collected in eight laboratory sites using two library preparation protocols (PolyA selection and rRNA depletion) and two RNA-seq devices (Illumina NovaSeq and MGI DNBSEQ-T7) in 21 batches^27^. Both alignment score and UMAP analysis have identified that all of these technical variations introduced unwanted variation into the transcriptomic data, with the variation caused by library preparation having the greatest impact (Figure 6a). Library preparation protocol variations are clearly visible in the UMAP analysis of the raw data (Figure 6b). Notably, these unknown variations, as well as batch variations, are greatly reduced in DeepAdapter-aligned data (Figure 6a-c & Figure S8). We systematically examine the transcriptomic profiles collected from different laboratories and devices but by the same protocol. The results reveal that the laboratory/device variations have persisted even after ruling out the major deviations brought about by the protocol, further confirming the complicated mixed bias introduced by multiple technical variations (Figure 6a & Figure S9). Interestingly, DeepAdpater adaptively learns from the dataset and efficiently eliminates the laboratory/device variations in this new situation (Figure S9). These results robustly illustrate DeepAdapter’s efficacy in simultaneously removing multiple mixed/unknown variations whose sources are challenging to identify.

**Figure 6.**
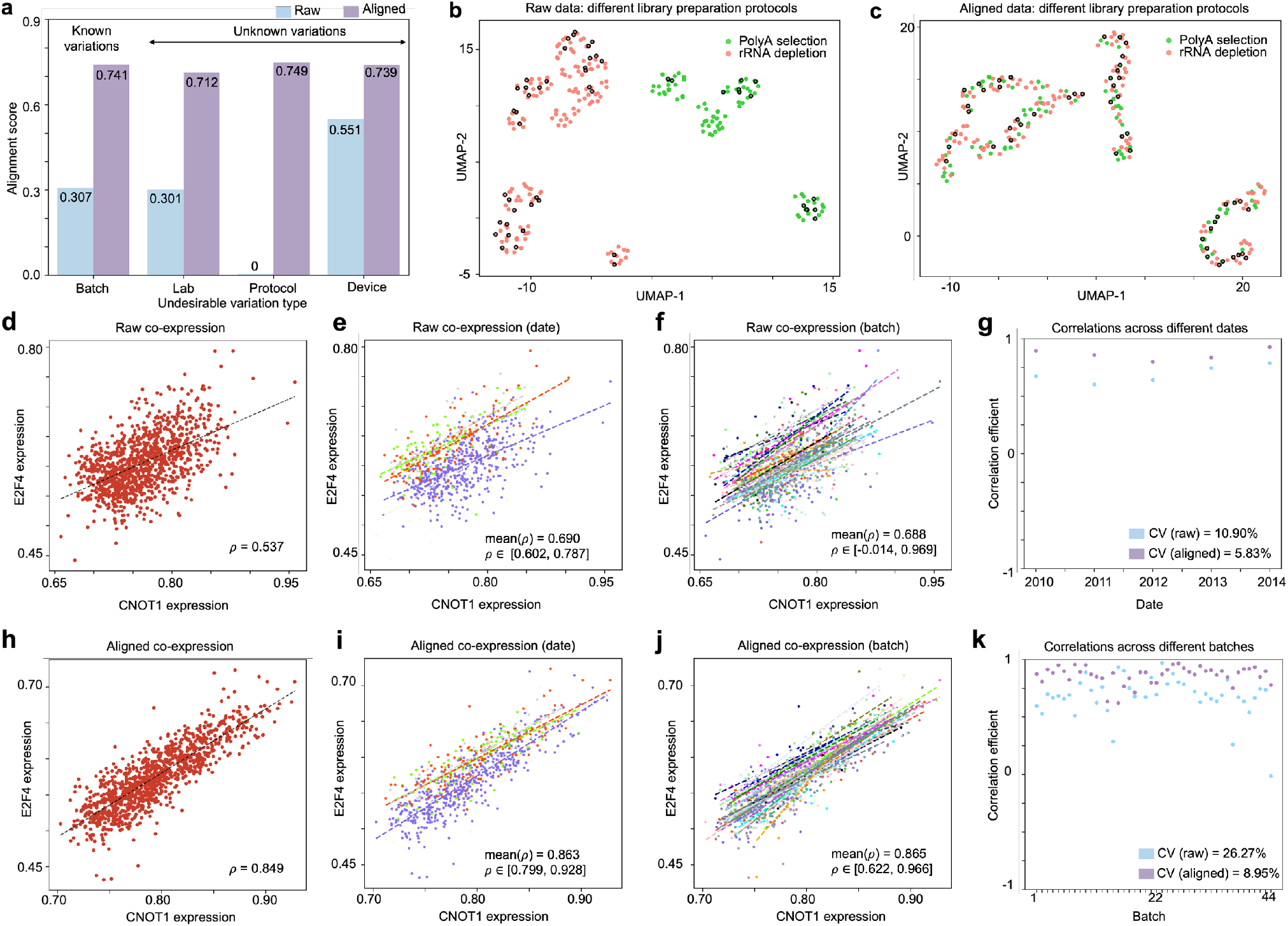
DeepAdapter corrects the unknown/mixed variations. a) The quantitative metrics of known (batch) and unknown (library protocols, laboratory sites, and devices) variations for Batch-Quartet data. The UMAP analysis of b) raw and c) aligned data across library preparation protocols. D) Correlation analysis for the expression level of CNOT1 and E2F4 in raw transcriptomic data. Correlation analysis and variation analysis of e) 5 processing dates and f) 44 experimental batches for CNOT1 and E2F4 gene expression in raw transcriptomic data. h) Correlation analysis for CNOT1 and E2F4 gene expression in aligned transcriptomic data. Correlation analysis and variation analysis of i) 5 processing dates and j) 44 experimental batches for the expression level of CNOT1 and E2F4 in aligned transcriptomic data. Coefficient of variations analysis for (g) dates and (i) batches.

#### DeepAdapter reproduces the relationship of gene co-expression with eliminated interferences in dates and batches

Undesirable variations can compromise gene co-expression analyses. For example, CNOT1 (CCR4-NOT transcription complex subunit 1) and E2F4 (E2F transcription factor 4) are demonstrated as the correlated gene pair for BRCA in both RNA-seq and microarray transcriptomic data^11^. Yet, the co-expression analysis between CNOT1 and E2F4 shows a limited correlation coefficient of 0.537 in unaligned TCGA data (Figure 6d). After DeepAdapater alignment, a strong correlation between the expression level of these two genes is detected (*ρ* =0.849, Figure 6h). Further analysis reveals that this is greatly affected by unwanted variation from complex sources: batches, processing dates and potentially other sources. By regrouping the raw data by processing dates and batches separately, the average correlation of these two genes within each group is largely improved (*ρ*=0.690/0.688 for 5 dates groups/44 batches groups, Figure 6e-g) compared to the overall correlation (*ρ*=0.537). After alignment, the regrouped data also shows a high average correlation (*ρ*=0.863/0.865 for dates/batches, Figure 6i-k). These results show that DeepAdapter corrects the unknown variations caused by complex mixed factors without prior knowledge.

## Discussion

In summary, we propose the first universal framework to eliminate the variations caused by a single factor or a combination of factors, including but not limited to well-known sources such as batches, platforms, tumor purities, etc. Utilizing transcriptomic datasets obtained from LINCS, Quartet, TCGA, GDSC, and CCLE, we reveal that DeepAdapter is efficient and robust in removing these variations and preserving meaningful biological variability for downstream analysis.

By carefully analyzing these large-scale transcriptomic data above, we illustrate the presence of numerous undesirable variations from different sources in the data, influencing downstream analysis and distorting biological insights. These variations tend not to exist in isolation, but are widely co-existed in the data, including from a single independent study and multiple different studies. In such situations, prevailing models that specialize in handling a specific type of undesirable variation^10^ are limited. A suitable model must be versatile in efficiently identifying and addressing different types of unwanted variations. We take advantage of the deep generative networks coupled with adversarial learning and contrastive learning to automatically correct multiple types of undesirable variations. Our results clearly show that DeepAdaptor can remove different sources of unwanted variations simultaneously. Therefore, we believe that it will be a valuable tool for biological scientists to better integrate and analyze large-scale transcriptomes.

The versatility of our model substantially advances the integrative investigations of the large and heterogeneous transcriptomes. Continuous innovation in RNA profiling technologies has resulted in the accumulation of extensive datasets. However, due to undesirable variations introduced by different technologies, they were not fully utilized. Our approach offers a practical solution to systematically eliminate these variations, enabling deep mining by integrating the historical datasets generated by early-developed technologies.

A remarkable feature of DeepAdapter is its capability in reconstructing transcriptomic profiles of pure tumor cells from bulk RNA data of highly heterogeneous tumor tissue by removing the purity variations. Although single-cell RNA-seq technology enables the detection of transcriptomic signatures of tumor cells, the cost has largely limited its clinical application. As an alternative, our work has demonstrated the feasibility of inferring the transcriptomic profiles of tumor cells from bulk RNA by eliminating the inevitable interferences of normal cells. We suggest that DeepAdapter could be used as a powerful tool for advancing clinical molecular diagnostics and precision medicine.

One limitation of our design is that DeepAdapter has been deployed to proofread the expression level of shared genes across different data, but not the whole transcriptome. Retaining only the intersecting genes sacrifices the integrity of the data. In the future, we plan to extend DeepAdapter’s applications and reconstruct the whole transcripts by revealing the latent relationship of the transcriptomic network.

## Data availability

All transcriptomic datasets in this study are publicly available. RNA-seq profiles of Batch-LINCS and Batch-Quartet can be downloaded from LINCS project (http://lincsportal.ccs.miami.edu/dcic-portal/) and Quartet (http://chinese-quartet.org) project. Microarray of cancer cell lines can be downloaded from GDSC project (https://www.cancerrxgene.org/). RNA-seq of cancer cell lines can be downloaded from CCLE project (https://depmap.org/portal/). RNA-seq of human tumor tissues can be downloaded from the TCGA project (https://www.cancer.gov/ccg/research/genome-sequencing/tcga).

## Code availability

The codes are available at https://github.com/mjDelta/DeepAdapter.

## Author Contributions

M. Z., T. X., B. C., and D. Y. conceived and designed the study. M. Z. developed DeepAdapter and made figures and documentation. M. Z. and D. Y. contributed to the experimental design. M. Z. performed experiments and statistical analysis. D. Y. provided insights into the application of the model. M. Z., Y. Y., S. Z., Y. Y., X. W., B. C. and D. Y. contributed to the data interpretation. M. Z. and S. Y. contributed to data acquisition. M. Z. and D.Y. wrote the manuscript. M. Z. and D. Y. reviewed and revised the manuscript.

## Acknowledgements

This study was funded by FosunLead, Inc.

## References

1. Cieślik, M. & Chinnaiyan, A.M. Cancer transcriptome profiling at the juncture of clinical translation. Nature Reviews Genetics 19, 93–109 (2018).

2. Lockhart, D.J. et al. Expression monitoring by hybridization to high-density oligonucleotide arrays. Nature biotechnology 14, 1675–1680 (1996).

3. Nagalakshmi, U. et al. The transcriptional landscape of the yeast genome defined by RNA sequencing. Science 320, 1344–1349 (2008).

4. Byron, S.A., Van Keuren-Jensen, K.R., Engelthaler, D.M., Carpten, J.D. & Craig, D.W. Translating RNA sequencing into clinical diagnostics: opportunities and challenges. Nature Reviews Genetics 17, 257–271 (2016).

5. Chang, J.C. et al. Gene expression profiling for the prediction of therapeutic response to docetaxel in patients with breast cancer. The Lancet 362, 362–369 (2003).

6. Robinson, D.R. et al. Integrative clinical genomics of metastatic cancer. Nature 548, 297–303 (2017).

7. Weinstein, J.N. et al. The cancer genome atlas pan-cancer analysis project. Nature genetics 45, 1113–1120 (2013).

8. Iorio, F. et al. A landscape of pharmacogenomic interactions in cancer. Cell 166, 740–754 (2016).

9. Ghandi, M. et al. Next-generation characterization of the Cancer Cell Line Encyclopedia. Nature 569, 503–508 (2019).

10. Haghverdi, L., Lun, A.T., Morgan, M.D. & Marioni, J.C. Batch effects in single-cell RNA-sequencing data are corrected by matching mutual nearest neighbors. Nature biotechnology 36, 421–427 (2018).

11. Molania, R. et al. Removing unwanted variation from large-scale RNA sequencing data with PRPS. Nature Biotechnology 41, 82–95 (2023).

12. Gagnon-Bartsch, J.A. & Speed, T.P. Using control genes to correct for unwanted variation in microarray data. Biostatistics 13, 539–552 (2012).

13. Domcke, S., Sinha, R., Levine, D.A., Sander, C. & Schultz, N. Evaluating cell lines as tumour models by comparison of genomic profiles. Nature communications 4, 2126 (2013).

14. Warren, A. et al. Global computational alignment of tumor and cell line transcriptional profiles. Nature Communications 12, 22 (2021).

15. Leek, J.T. et al. Tackling the widespread and critical impact of batch effects in high-throughput data. Nature Reviews Genetics 11, 733–739 (2010).

16. Sims, D., Sudbery, I., Ilott, N.E., Heger, A. & Ponting, C.P. Sequencing depth and coverage: key considerations in genomic analyses. Nature Reviews Genetics 15, 121–132 (2014).

17. Wang, C. et al. The concordance between RNA-seq and microarray data depends on chemical treatment and transcript abundance. Nature Biotechnology 32, 926–932 (2014).

18. Johnson, W.E., Li, C. & Rabinovic, A. Adjusting batch effects in microarray expression data using empirical Bayes methods. Biostatistics 8, 118–127 (2007).

19. Zhang, Y., Parmigiani, G. & Johnson, W.E. ComBat-seq: batch effect adjustment for RNA-seq count data. NAR genomics and bioinformatics 2, lqaa078 (2020).

20. Büttner, M., Miao, Z., Wolf, F.A., Teichmann, S.A. & Theis, F.J. A test metric for assessing single-cell RNA-seq batch correction. Nature methods 16, 43–49 (2019).

21. Ledford, H. The death of microarrays? Nature 455, 847 (2008).

22. Fu, X. et al. Estimating accuracy of RNA-Seq and microarrays with proteomics. BMC genomics 10, 1–9 (2009).

23. Dillies, M.-A. et al. A comprehensive evaluation of normalization methods for Illumina high-throughput RNA sequencing data analysis. Briefings in bioinformatics 14, 671–683 (2013).

24. Bullard, J.H., Purdom, E., Hansen, K.D. & Dudoit, S. Evaluation of statistical methods for normalization and differential expression in mRNA-Seq experiments. BMC bioinformatics 11, 1–13 (2010).

25. Newman, A.M. et al. Robust enumeration of cell subsets from tissue expression profiles. Nature Methods 12, 453–457 (2015).

26. van Hasselt, J.G.C. et al. Transcriptomic profiling of human cardiac cells predicts protein kinase inhibitor-associated cardiotoxicity. Nature Communications 11, 4809 (2020).

27. Yu, Y. et al. Quartet RNA reference materials improve the quality of transcriptomic data through ratio-based profiling. Nature Biotechnology (2023).

28. Stark, R., Grzelak, M. & Hadfield, J. RNA sequencing: the teenage years. Nature Reviews Genetics 20, 631–656 (2019).

29. Goodfellow, I.J. et al./person-group>. in Advances in Neural Information Processing Systems 27, Vol. 27. (eds. Z. Ghahramani, M. Welling, C. Cortes, N.D. Lawrence & K.Q. Weinberger) (2014).

30. Zhao, Y., Wong, L. & Goh, W.W.B. How to do quantile normalization correctly for gene expression data analyses. Scientific reports 10, 15534 (2020).

31. McInnes, L., Healy, J. & Melville, J. Umap: Uniform manifold approximation and projection for dimension reduction. arXiv preprint arXiv:1802.03426 (2018).

32. Butler, A., Hoffman, P., Smibert, P., Papalexi, E. & Satija, R. Integrating single-cell transcriptomic data across different conditions, technologies, and species. Nature biotechnology 36, 411–420 (2018).

33. Pedregosa, F. et al. Scikit-learn: Machine learning in Python. the Journal of machine Learning research 12, 2825–2830 (2011).

34. Hubert, L. & Arabie, P. Comparing partitions. Journal of classification 2, 193–218 (1985).

35. Hanahan, D. & Weinberg, R.A. Hallmarks of cancer: the next generation. cell 144, 646–674 (2011).

36. Chu, T., Wang, Z., Pe’er, D. & Danko, C.G. Cell type and gene expression deconvolution with BayesPrism enables Bayesian integrative analysis across bulk and single-cell RNA sequencing in oncology. Nature Cancer 3, 505–517 (2022).

37. Yoshihara, K. et al. Inferring tumour purity and stromal and immune cell admixture from expression data. Nature Communications 4, 2612 (2013).

38. Liu, Z. et al. A glycolysis-related two-gene risk model that can effectively predict the prognosis of patients with rectal cancer. Human Genomics 16, 5 (2022).

