## supplementary information for "A self-adaptive and versatile tool for eliminating multiple undesirable variations from transcriptome"

#### Title

#### Methods

##### Datasets

***Transcriptomic data:*** The transcriptomic data includes gene expression datasets across batches, sequencing platforms and bio-samples. For batch variations, RNA-seq profiles of cell lines are sourced from LINCS project (http://lincsportal.ccs.miami.edu/dcic-portal/) and Quartet (http://chinese-quartet.org) project. For platform variations, microarray of cell lines is obtained from The Genomics of Drug Sensitivity in Cancer (GDSC) project (https://www.cancerrxgene.org). RNA-seq of cell lines comes from DepMap (22Q2 project, https://depmap.org/portal/). For purity variations, RNA-seq of cell lines comes from DepMap (19Q4 project, https://depmap.org/portal/), and RNA-seq of tumor tissues comes from The Cancer Genome Atlas (TCGA) project[1] (https://www.cancer.gov/ccg/research/genome-sequencing/tcga).

***Survival data:*** The survival data of the TCGA project are downloaded from the GDC Data Portal (https://portal.gdc.cancer.gov/).

***Batches and processing dates annotations:*** The annotations of batches and processing dates for TCGA project are downloaded from the MBatch Omic Browser (https://bioinformatics.mdanderson.org/MQA/).

***Known potential prognostic genes***: The genes with significance in overall survival analysis are obtained from two sources: 1) the presented results of PRPS project[2], which aims to remove the variations brought by tumor purity and batch effects for TCGA project; 2) targeted research on glycolysis-related gene sets in prognosis analysis for colon adenocarcinoma (COAD) and rectal adenocarcinoma (READ)[3]. Specifically, there exists 8, 6, and 3 known potential prognostic genes for COAD, READ, and BRCA, respectively: 1) SUCLG2P2, GPC1, CISH, ANKZF1, BRAF, P4HA1, STC2, and PCK1 for COAD; 2) TSTA3, RAB18, PTPN14, PKP2, FBXL14, and CSGALNACT2 for READ; 3) ZEB2, STAB1, and ESRRA for BRCA.

##### The architecture of DeepAligner

DeepAligner is designed in an autoencoder structure and the unwanted variations are removed in the latent space between encoder and decoder. The encoder $\boldsymbol{E}$ takes the gene expression vector as the input $\boldsymbol{x}$ and extracts the latent vector with fully connected layers. With the extracted latent vector $\boldsymbol{l}$ projected into a low-dimensional space, DeepAligner corrects the artifacts with adversarial learning and deep metric learning. After removing the unwanted variations, the decoder $\boldsymbol{D}$ reconstructs the transcriptomic data $\boldsymbol{x'}$ by projecting the latent vector back into the original high-dimensional space.

The adversarial learning removes the artifacts by introducing a discriminator $\boldsymbol{F}$, which is designed to recognize the labels of sequencing platforms and biosamples. During adversarial learning, the encoder $\boldsymbol{E}$ is trained to confuse the discriminator $\boldsymbol{F}$ in a min-max optimization process, thereby correcting the unwanted variations across sequencing platforms and biosamples.

The deep metric learning removes the artifacts by minimizing the distance between the anchor and the positive and maximizing the distance between the anchor and the negative. DeepAligner automatically defines the positive samples with mutual nearest neighbors[4] without any additional biological annotations as follows:

$$(m,n) \in Space_{a,b} \text{ and } (n,m) \in Space_{a,b} \Leftrightarrow \begin{cases} m \in Source_a, n \in Source_b \\ m \in \mathrm{NN}(x_n^b, X^a) \text{ and } n \in \mathrm{NN}(x_m^a, X^b) \end{cases}, \quad (1)$$

where $\boldsymbol{m}$ and $\boldsymbol{n}$ refer to the extracted MNN pair, $\mathrm{NN}(x_n^b, X^a)$ refers to the nearest neighbors in source $\boldsymbol{a}$ of sample $\boldsymbol{n}$ from source $\boldsymbol{b}$, $\mathrm{NN}(x_m^a, X^b)$ refers to the nearest neighbors in source $\boldsymbol{b}$ of sample $\boldsymbol{m}$ from source $\boldsymbol{a}$. The negative is defined as the randomly selected samples from the same source, excluded the sample itself.

**Loss function:**

The loss function includes 3 parts: the reconstruction loss of autoencoder, the adversarial loss of the discriminator, and the triplet loss of the deep metric learning. The reconstruction loss is defined with mean absolute error as follows:

$$\mathrm{loss}_{rec} = \sum_{x_i \in x, x_i' \in x'} |x_i - x_i'|, \quad (2)$$

where $\boldsymbol{x}$ and $\boldsymbol{x'}$ refer to the original and reconstructed transcriptomic data. The classification loss of the discriminator $\boldsymbol{F}$ is defined with cross-entropy as follows:

$$\mathrm{loss}_{\mathrm{disc}} = CE(\mathrm{F}(l), y^{\mathrm{s}}), \quad (3)$$

where $\boldsymbol{l}$ refers to the latent vector extracted by encoder $\boldsymbol{E}$, $\boldsymbol{y^s}$ refers to the source label (e.g., RNA-seq/microarray for different sequencing platforms or cell line/tumor tissue for different biosamples) of transcriptomic data, and *CE* refers to the cross-entropy calculation. The triplet loss is defined as follows:

$$\mathrm{loss}_{\mathrm{triplet}} = max(d(a,p) - d(a,n) + m, 0), \quad (4)$$

where $\boldsymbol{a}$, $\boldsymbol{p}$, $\boldsymbol{n}$ refer to the anchor, the positive, and the negative sample defined in mutual nearest neighbors, $\boldsymbol{d}$ refers to the distance metric measured by Euclidean distance, and $\boldsymbol{m}$ refers to the margin value of 1.0. With reconstruction loss, classification loss, and triplet loss, the total loss function of DeepAdapter is defined with min-max optimization as follows:

$$\min_{E,D} \max_{F} L(E,D,F) = loss_{rec} + \mathrm{loss}_{\mathrm{triplet}} - \lambda \mathrm{loss}_{\mathrm{disc}}, \quad (5)$$

where $\lambda = 0.01$ refers to the weight of adversarial learning.

#### Training of DeepAdapter

DeepAdapter is trained with $epochs = 150K$ using Adam optimizer with $\beta_1 = 0.9$ and $\beta_2 = 0.98$. The learning rate is first increased linearly to the maximum learning rate of $5 \times 10^{-4}$ and then decreased linearly to the minimum learning rate of $10^{-5}$. The batch size is set as 256. The whole model is trained with single Nvidia GeForce RTX 3090 Ti.

#### Evaluation metrics

Two sets of evaluation metrics are included in this work: 1) variation removal and 2) biological signal conservation.

Variation removal metrics include alignment score[5] and modified average silhouette width (ASW) score[6]. The basic intuition of variation removal is that if the samples are well aligned, the neighbors of any sample are evenly distributed across data sources. Alignment score measures the distribution of samples from different sources around one sample. The alignment score is calculated as follows:

$$\text{alignment score} = 1 - \frac{\bar{x} - \frac{k}{N}}{k - \frac{k}{N}}, \tag{6}$$

where $\boldsymbol{k}$ refers to the number of nearest neighbors, $\boldsymbol{N}$ refers to the total number of samples, $\bar{x}$ refers to the average number of nearest neighbors belonging to the same data source. The original ASW score is the average value of all silhouette widths of all samples and is defined as follows:

$$\text{original ASW} = \sum\nolimits_{i \in N} silhouette\ width(i), \tag{7}$$

where i refers to the sample i and the silhouette width measures the similarity consistency within clusters:

$$silhouette\ width(i) = \frac{(b - a)}{\max(a, b)}, \tag{8}$$

where a and b refer to the intra-cluster and nearest-cluster distance for sample i. Thus, the modified ASW for assessing variation correction measures the opposite aspect of within-cluster consistency and is defined as follows:

$$\text{modified ASW} = 1 - \text{original ASW}. \tag{9}$$

Biological signal conservation metrics include normalized mutual information (NMI)[7] and adjusted rand index (ARI)[8]. Both NMI and ARI compare the overlap between biological annotations and clustered annotations calculated from the integrated dataset. NMI is calculated with entropy as follows:

$$NMI = \frac{2 \times I(Y; C)}{[\mathrm{H(Y)} + \mathrm{H(C)}]}, \tag{10}$$

where Y and C refer to the biological and clustered annotations, H(Y) and H(C) refer to the entropy of biological and clustered annotations, and I(Y;C) refers to the mutual information between biological and clustered annotations. ARI is calculated as follows:

$$ARI = \frac{\sum_{ij}\binom{n_{Y_iC_j}}{2} - \sum_i\binom{n_{Y_i}}{2}\sum_j\binom{n_{C_j}}{2}/\binom{n}{2}}{\frac{1}{2}\left(\sum_i\binom{n_{Y_i}}{2} + \sum_j\binom{n_{C_j}}{2}\right) - \sum_i\binom{n_{Y_i}}{2}\sum_j\binom{n_{C_j}}{2}/\binom{n}{2}}, \tag{11}$$

where $n_{Y_iC_j}$ refers to the number of samples of the biological annotation $Y_i$ assigned to cluster $C_j$, $n_{Y_i}$ refers to the number of samples in biological annotation $Y_i$, and $n_{C_j}$ refers to the number of samples in the cluster annotation $C_j$.

Jensen-Shannon divergence (JSD) is used to measure the similarity of gene expression distributions for matched RNA-seq and microarray of cell lines. Jensen-Shannon divergence is calculated as follows:

$$JSD = \frac{1}{2}D(P \parallel M) + \frac{1}{2}D(Q \parallel M), \tag{12}$$

where $M = 1/2(P + Q)$, and P and Q refer to the gene expression distributions for matched RNA-seq and microarray of cancer cell lines, respectively.

#### Baseline approaches

Quantile normalization is implemented with qnorm (version 0.8.1). Combat[9] corrects the unwanted variations with an empirical Bayes design and is implemented with pycombat (version 0.3.3, https://epigenelabs.github.io/pyComBat/). MNN[4] corrects the bias with the correction vectors between MNN pairs and is implemented with mnnpy (version 0.1.9.5, https://pypi.org/project/mnnpy/).

#### Tumor purity calculation

The inferred tumor purity of bulk gene expression profiles is calculated with ESTIMATE by inferring the proportion of stromal and immune cells[10].

#### Statistical analysis

Calculation of Pearson correlation coefficients is implemented with scipy (version 1.11.1, www.scipy.org). UMAP decomposition is implemented with umap-learn (version 0.5.3, https://umap-learn.readthedocs.io). AWS, NMI, and ARI is implemented with scikit-learn (version 1.2.2, www.scikit-learn.org). Kaplan Meier analysis is implemented with lifelines (version 0.27.7, https://lifelines.readthedocs.io). Log rank test for survival analysis is implemented with scikit-survival (version 0.21.0, https://scikit-survival.readthedocs.io/en/stable/).

### Figures

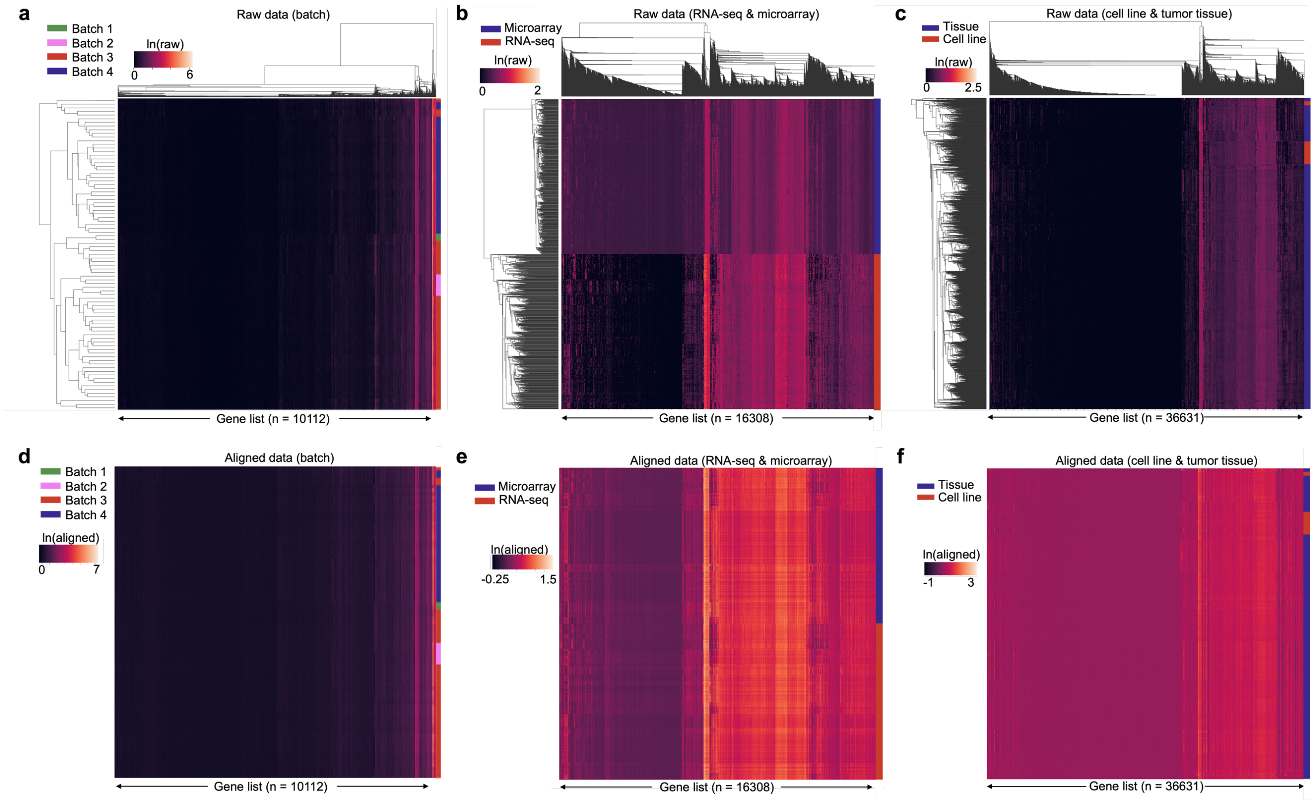


Figure S1. Heatmap of transcriptomic profiles. Raw data for different a) batches, b) platforms, and c) bio-samples. Aligned data for different d) batches, e) platforms, and f) bio-samples.

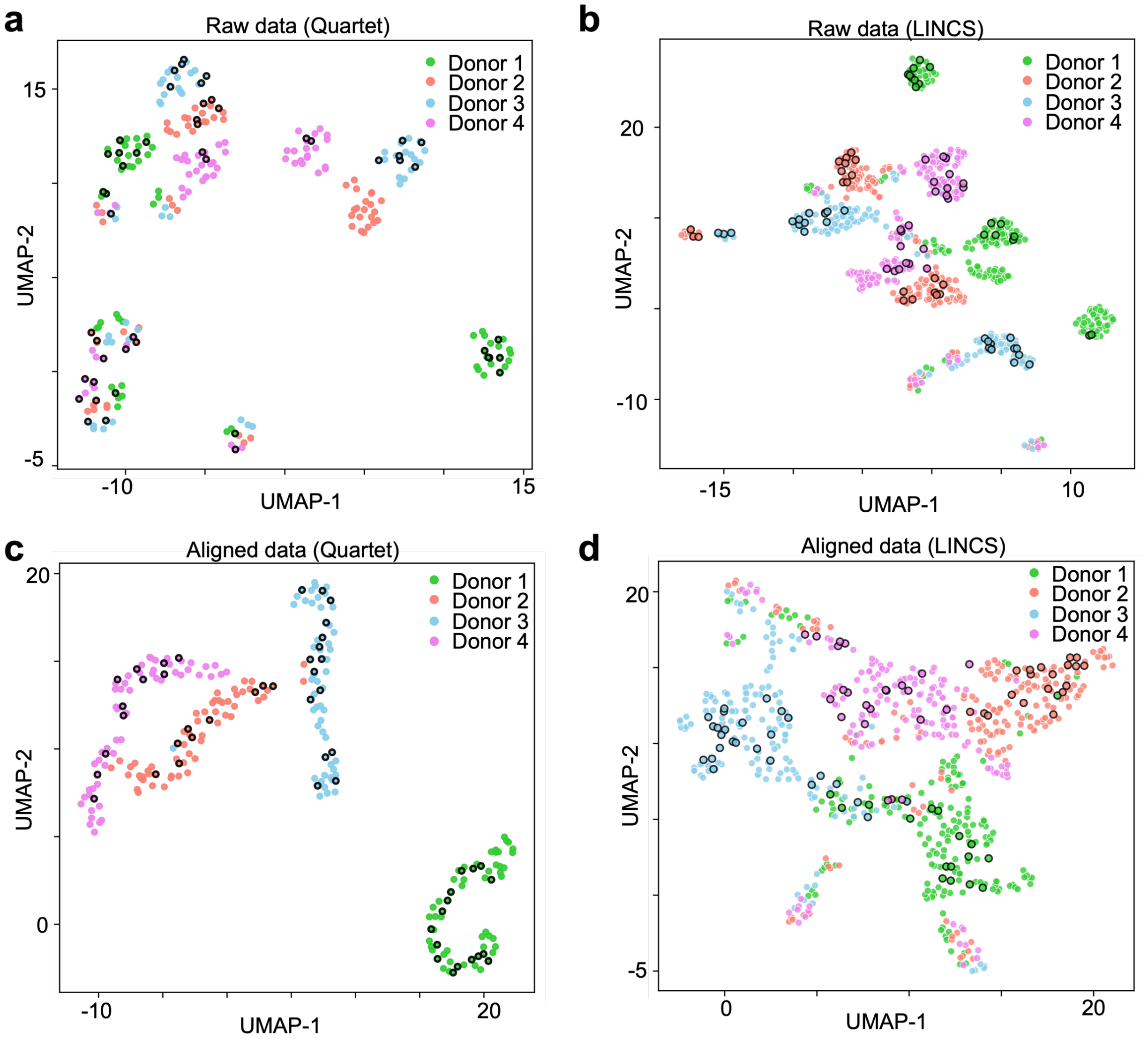


Figure S2. UMAP analysis of the distinct donor-wise signals. UMAP analysis of raw data for a) Batch-Quartet and b) Batch-LINCS. UMAP analysis of aligned data for c) Batch-Quartet and d) Batch-LINCS.

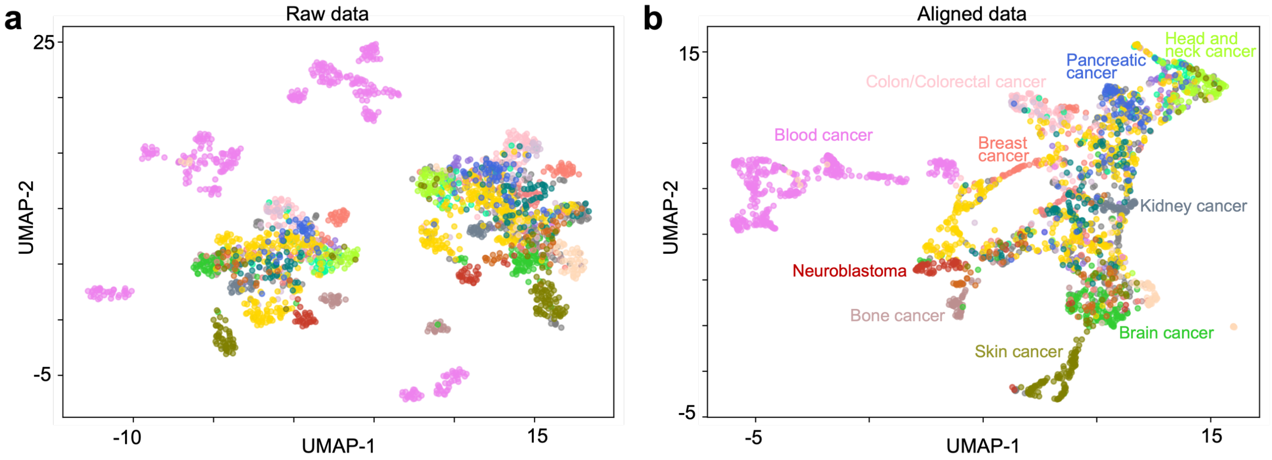


Figure S3. Overview of cancer type signals for platform variations. UMAP analysis of a) raw and b) aligned data across microarray and RNA-seq technologies.

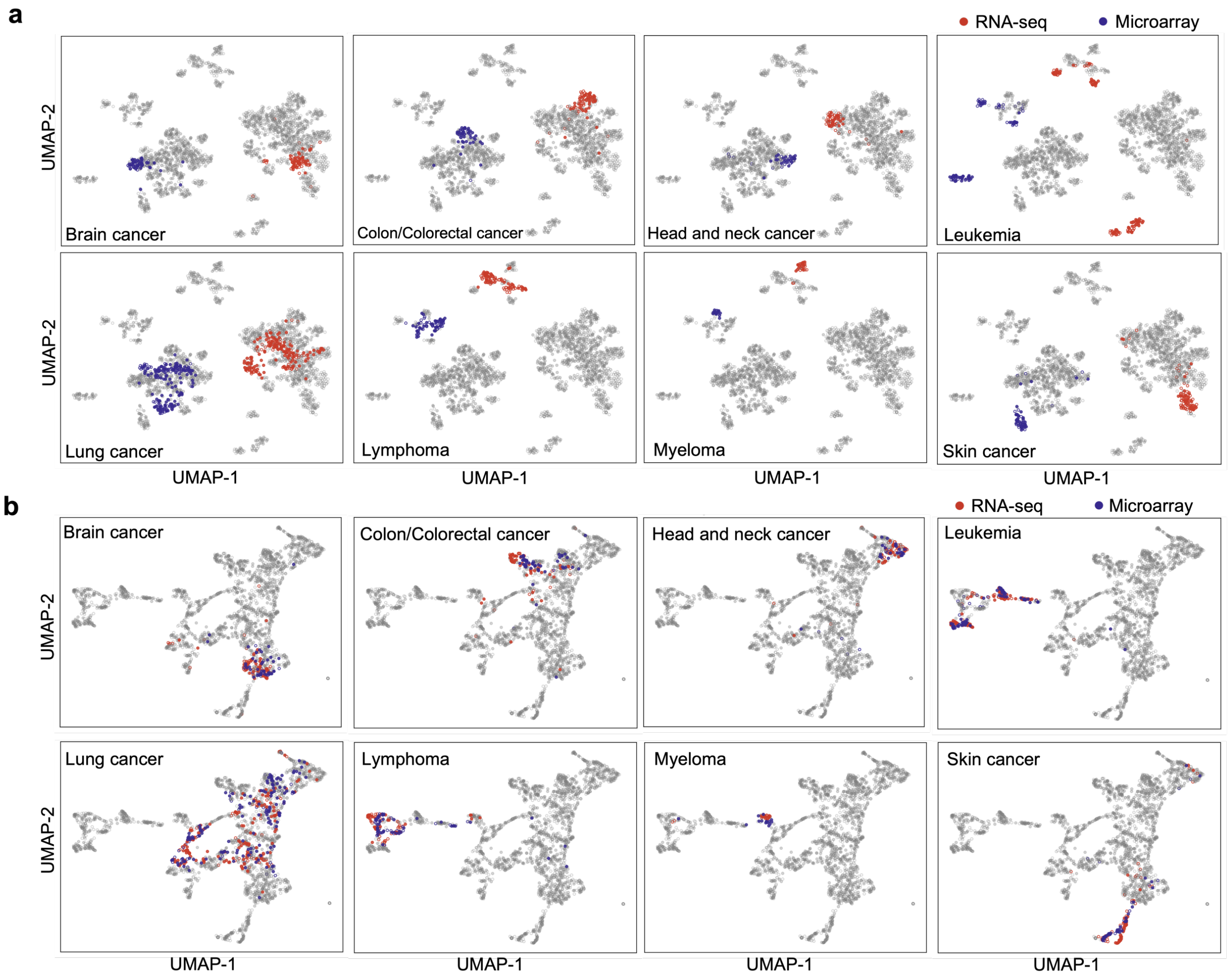


Figure S4. UMAP analysis of cell lines profiled by RNA-seq and microarray across cancer types. a) Separated profiles of RNA-seq and microarray across cancer types for raw transcriptomic data. b) Mixed profiles of RNA-seq and microarray across cancer types for aligned transcriptomic data. Overlapped microarray and RNA-seq samples indicates the removal of platform variations. Empty and filled circles refer to the training and testing set, respectively.

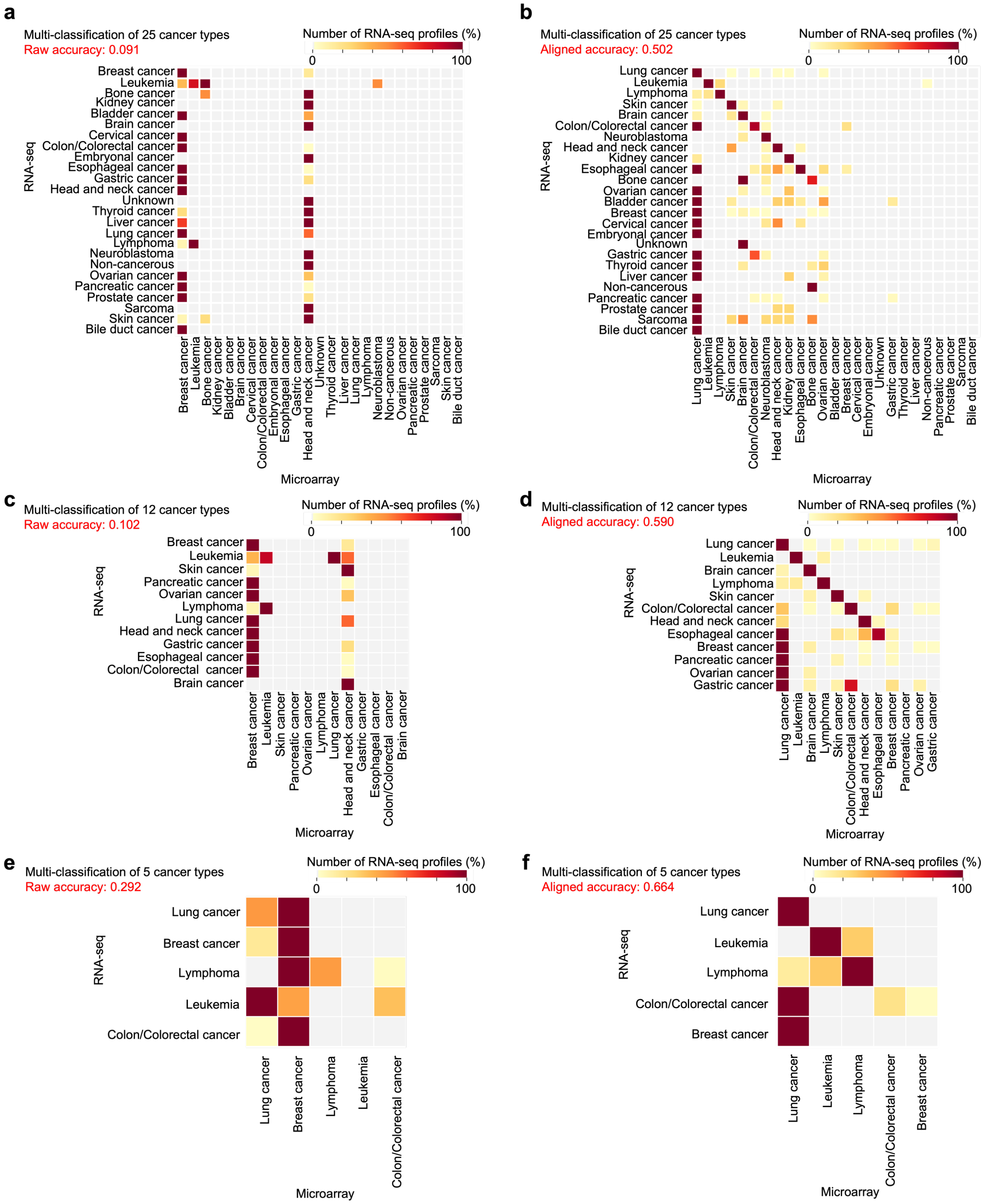


Figure S5. Confusion matrix of cross-platform multiclassification for cancer cell lines. Using the raw data, multiclassification of a) 25, c) 12, and e) 5 cancer types. Using the aligned data, multiclassification of b) 25, d) 12, and f) 5 cancer types.

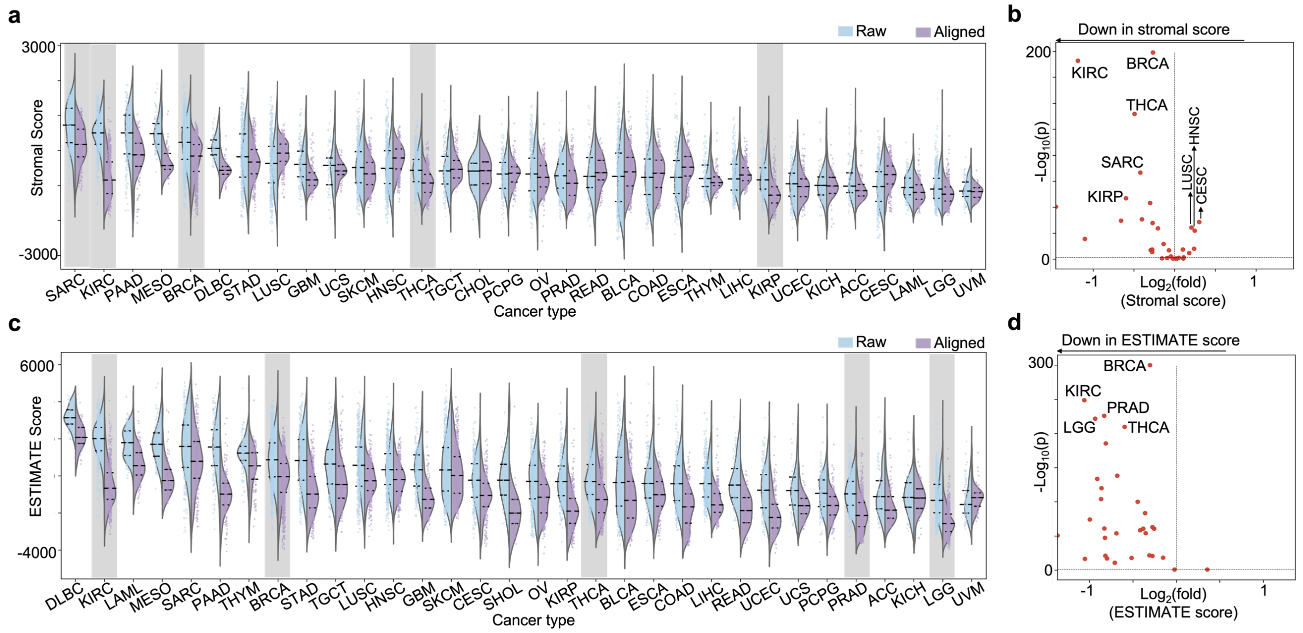


Figure S6. ESTIMATE analysis for transcriptomic profiles. a) Distribution of stromal score for raw and aligned data. b) Volcano plot showing the altered stromal score. c) Distribution of ESTIMATE score for raw and aligned data. d) Volcano plot showing the altered ESTIMATE score.

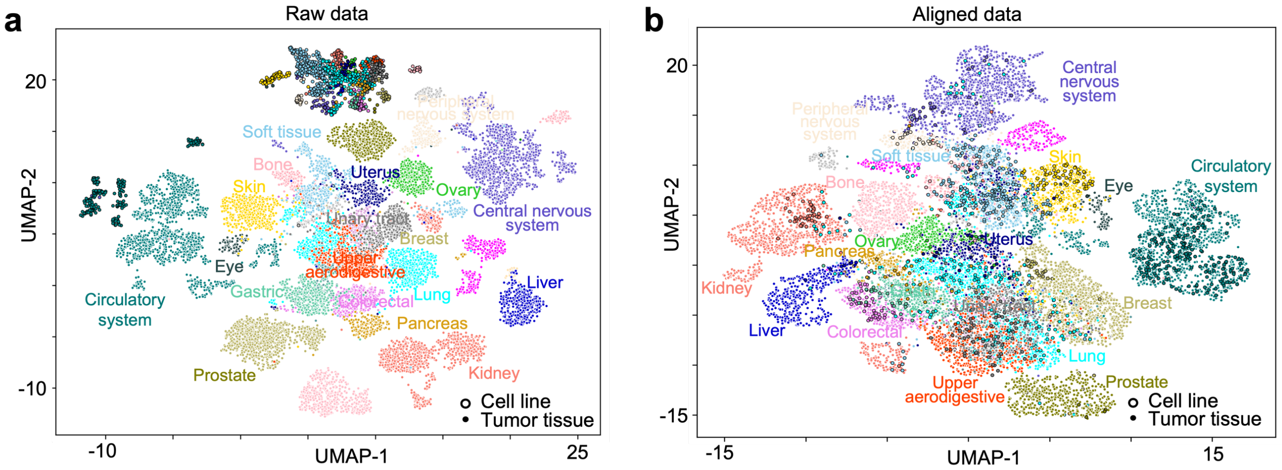


Figure S7. Overview of lineage distribution for purity variations. UMAP 2D projection of a) raw and aligned data across cancer cell lines and tumor tissues. Overlapped cell lines and tumor tissues indicates the removal of purity variations. Colors refer to cancer types. Circles with black and white edges refer to the data profiled from cell lines and tissues, respectively.

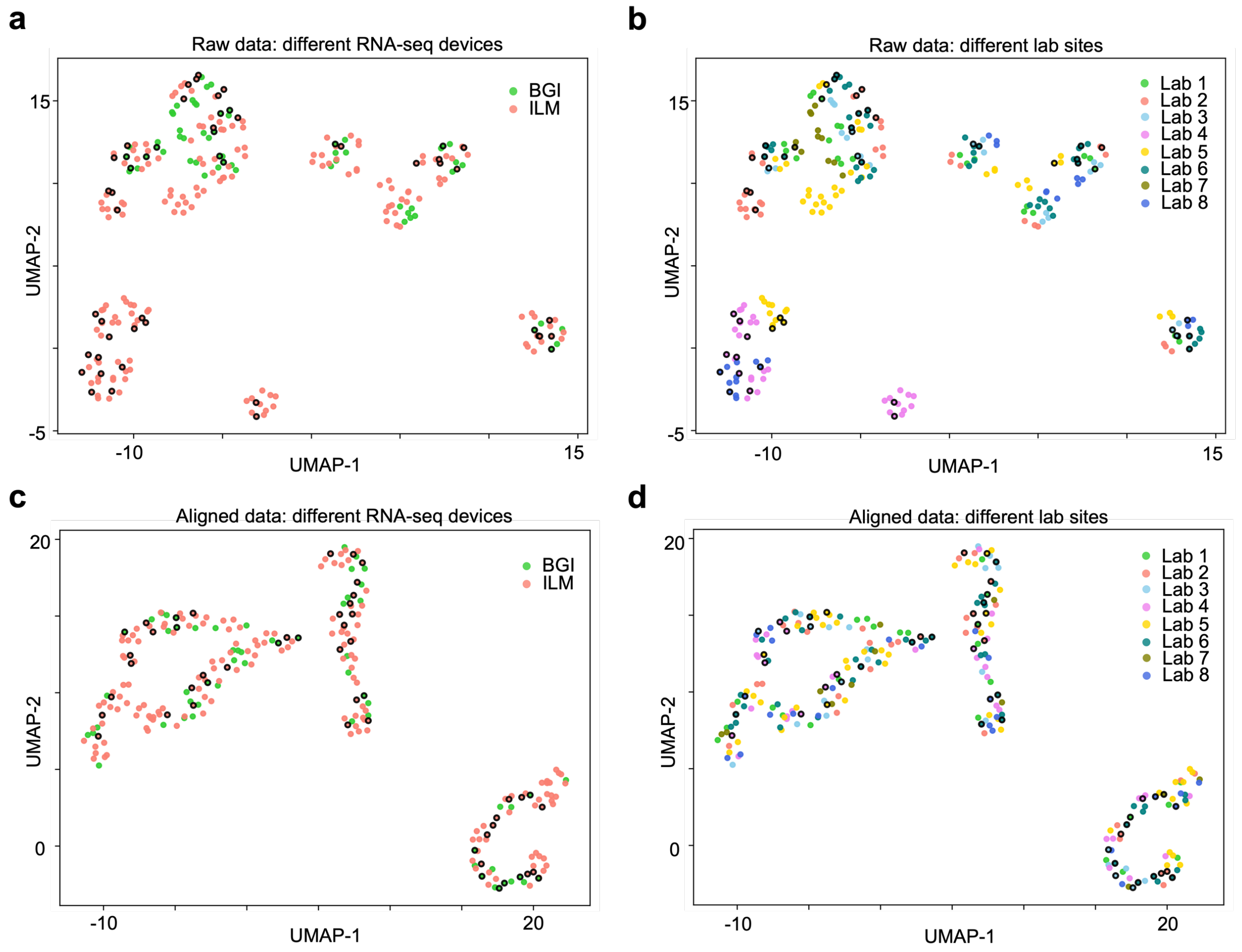


Figure S8. UMAP analysis of unknown/mixed variations for Batch-Quartet. The UMAP analysis of raw data across a) RAN-seq devices and b) laboratories. The UMAP analysis of aligned data across c) RAN-seq devices and d) laboratories.

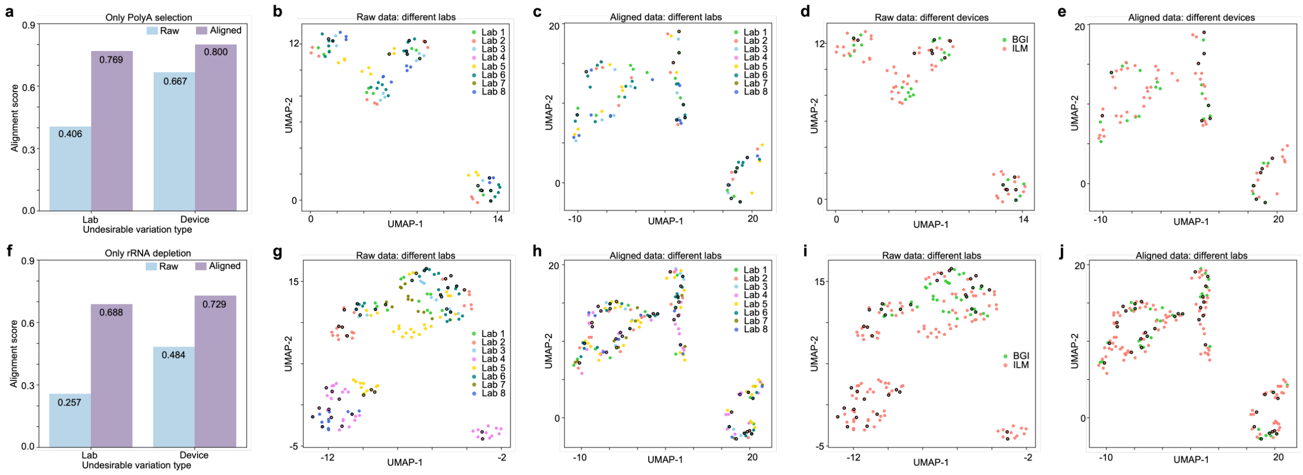


Figure S9. UMAP analysis of different laboratories and RNA-seq devices within the same library preparation protocols. The quantitative metrics of laboratory sites and devices (unknown) variations within the a) polyA selection and f) rRNA depletion protocol. The UMAP analysis of b) raw and c) aligned data across laboratories within polyA selection protocol. The UMAP analysis of d) raw and e) aligned data across RNA-seq devices within polyA selection protocol. The UMAP analysis of g) raw and h) aligned data across laboratories within rRNA depletion protocol. The UMAP analysis of i) raw and j) aligned data across RNA-seq devices within rRNA depletion protocol. White- and black-edged circles refer to the training and testing set, respectively.

#### References


1. Weinstein, J.N. et al. The cancer genome atlas pan-cancer analysis project. *Nature genetics* **45**, 1113-1120 (2013).
2. Molania, R. et al. Removing unwanted variation from large-scale RNA sequencing data with PRPS. *Nature Biotechnology* **41**, 82-95 (2023).
3. Liu, Z. et al. A glycolysis-related two-gene risk model that can effectively predict the prognosis of patients with rectal cancer. *Human Genomics* **16**, 5 (2022).
4. Haghverdi, L., Lun, A.T., Morgan, M.D. & Marioni, J.C. Batch effects in single-cell RNA-sequencing data are corrected by matching mutual nearest neighbors. *Nature biotechnology* **36**, 421-427 (2018).
5. Butler, A., Hoffman, P., Smibert, P., Papalexi, E. & Satija, R. Integrating single-cell transcriptomic data across different conditions, technologies, and species. *Nature biotechnology* **36**, 411-420 (2018).
6. Büttner, M., Miao, Z., Wolf, F.A., Teichmann, S.A. & Theis, F.J. A test metric for assessing single-cell RNA-seq batch correction. *Nature methods* **16**, 43-49 (2019).
7. Pedregosa, F. et al. Scikit-learn: Machine learning in Python. *the Journal of machine Learning research* **12**, 2825-2830 (2011).
8. Hubert, L. & Arabie, P. Comparing partitions. *Journal of classification* **2**, 193-218 (1985).
9. Johnson, W.E., Li, C. & Rabinovic, A. Adjusting batch effects in microarray expression data using empirical Bayes methods. *Biostatistics* **8**, 118-127 (2007).
10. Yoshihara, K. et al. Inferring tumour purity and stromal and immune cell admixture from expression data. *Nature Communications* **4**, 2612 (2013).
